## Supplementary material for "The evolution of antibiotic resistance islands occurs within the framework of plasmid lineages": Suplamentary Material

Supplementary Note

**Supplementary Note SN1. Demonstrative examples of the analysis pipeline outcome.**

This section presents two demonstrative examples of the analysis pipeline outcome (the pipeline is illustrated in Fig. 1A). As the first example, we show details on the identification of CSBs that correspond to Tn*125*[1]. This transposon is captured in our pipeline as it contains two cooccurring ARGs: *bla*$_{\text{NDM-1}}$ and *ble*$_{\text{MBL}}$ (Fig. SN1.1). The two gene families *bla*$_{\text{NDM-1}}$ and *ble*$_{\text{MBL}}$ were found to be significantly cooccurring in plasmids in the all three genera. Thus, these two ARGs form a coARG by our definition (all coARGs are listed in Supplementary Table S5). As the next step, we searched for CSBs that include both of the gene families in this coARG (i.e., *bla*$_{\text{NDM-1}}$ and *ble*$_{\text{MBL}}$). A total of 443 relevant CSBs were identified. These CSBs were searched against the transposable elements databases using sequence similarity. The six example CSBs containing this coARG were found to match Tn*125* (details in Supplementary Table S6). The counting of ARGs in resistance islands included all ARGs that are found in such 'pieces of resistance islands' (i.e., CSBs that include coARGs and match to known transposable elements).

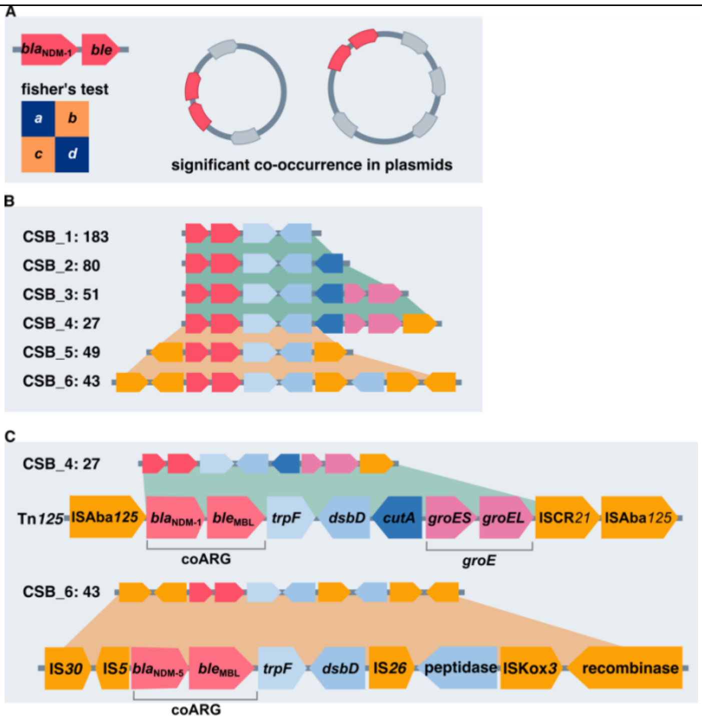

**Figure SN1.1. Example for piece-wise detection of Tn*125*.** (**A**) The genes *bla*$_{\text{NDM-1}}$ and *ble*$_{\text{MBL}}$ cooccur in plasmids more frequently than the expected by chance alone. (**B**) The two genes are found in six colinear syntenic blocks (CSBs). The CSBs are shown as arrows that correspond to specific genes in Tn*125* (shown below). The number of CSB instances (i.e., how many times we observed the CSB in plasmids) is shown after the CSB name. (**C**) CSB_3 and CSB_4 correspond to partial sequences of Tn*125*, with CSB_4 being the longest match to that transposon. CSB_5 and CSB_6 correspond to partial sequences of another Tn*125*-like sequence, where we find mostly *bla*$_{\text{NDM-5}}$ instead of *bla*$_{\text{NDM-1}}$ at the same place in Tn*125*. Note that the annotation of *dsbD* and *cutA* as a gene or pseudogene may depend on the version of RefSeq database. All the CSBs shown here are considered in our analysis as pieces of resistance islands.

Our gene family clustering approach relies on global sequence similarity hence it may cluster different ARG variants into the same gene family. This aspect of our analysis may lead to a bias in the reported coARGs (e.g., in the network analysis in Fig. 2A). Since the coARGs data is integrated with CSBs, such bias has a smaller effect in the CSBs final analysis outcome (e.g., Fig. 5C). This is because ARG variants that are associated with different mobile genetic elements will be distinguished in the CSB analysis. One example is the homologous genes $bla_{\text{NDM-1}}$ and $bla_{\text{NDM-5}}$ that are distinguished by the different CSBs in which they appear (Figure SN1.1). A second example is ARG family *dfrA*, which includes (in our data) two ARG variants *dfrA14* and *dfrA17*. The two variants were found in different CSBs in our analysis (Figure SN1.2). The CSB_*dfrA17* has coARGs but CSB_*dfrA14* carries single ARG, thus the latter was not included in our plasmid resistance island pipeline.

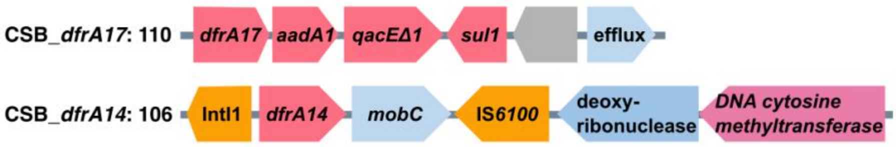

**Figure SN1.2. Example of ARG variants found in different CSBs**. ARGs *dfrA14* and *dfrA17* were clustered in the same gene family, but they are enriched in two different CSBs. CSB_*dfrA14* was excluded due to lack of coARG. The number of instances is shown after each CSB name. Protein-coding genes are shown as arrows, hypothetical gene is coloured in grey.

The CSBs we identified can be nested, as the example CSBs shown in Figure ST1.1. Thus, our approach does not distinguish the exact boundaries of smaller MGEs carrying ARGs (e.g., transposable elements). Alternatively, we consider the CSBs as 'pieces of resistance islands'. The location of ARGs and SSRs in CSBs was determined using the genomic coordinates of each gene. The size distribution of CSBs presented in Supplementary Figure S1 shows that the detected CSBs include not only intact MGEs carrying ARGs, but also partial MGE sequences (shorter CSBs), as well as MGE agglomerants (longer CSBs). An example for such an MGE agglomeration that is captured in our pipeline is demonstrated in Supplementary Figure S2.

### Supplementary Note SN2. Small and large *KES* plasmids are evolutionary diverged

Since plasmid mobility type and plasmid size are associated (i.e., conjugative plasmids are typically large), statistical statements on various association of plasmid gene content with plasmid mobility class alone may bias the process of conclusion making. Specifically in the context of MDR plasmids, if the positive association between plasmid size and number of ARGs is trivial, i.e., ARG acquisition leads to plasmid genome expansion, then large and small MDR plasmids are expected to be related (at least in some cases). In other words, large and small plasmids may have a shared gene content, with large plasmids harboring more ARGs in comparison to small plasmids. To test this hypothesis, one has to first find an objective approach to classify our plasmids into 'large' and 'small' plasmid groups. The distribution of plasmid size has been frequently described as a bi-modal distribution (as previously shown[2]). A similar bi-modal plasmid size distribution is observed in our data, with a minimal density at plasmid size of ca. 19kbp (Fig. SN2.1).

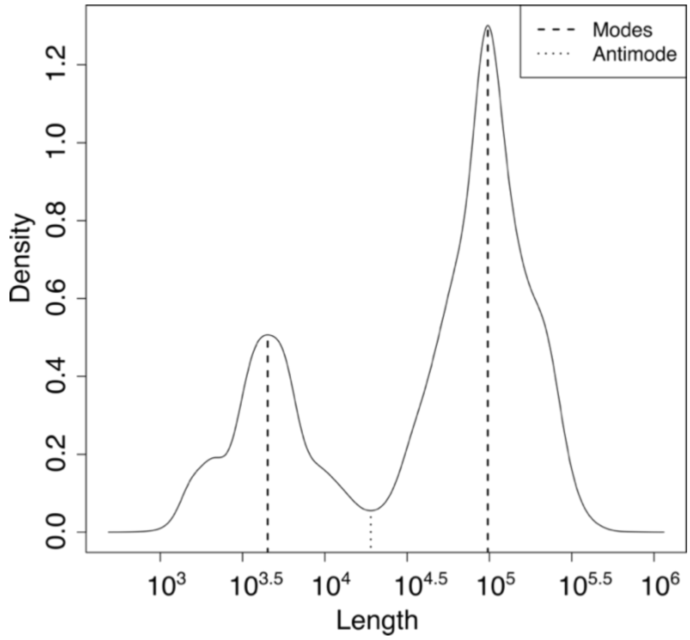

**Figure SN2.1. Density plot of genome size for all plasmids in the dataset.** Plasmid size is presented in a common logarithm scale. A bi-modal distribution is given for the estimation of the two highest density values (marked out with thick dash lines) and the lowest density value (marked out with a thin dash line). The multimodality of plasmid size distribution was tested using R function modetest. The minimum and maximum density of plasmid size distribution were estimated using R function locmodes with a bimodal distribution.

Using the minimum density value as a threshold, we compared the gene content between small plasmids having a genome size <19 kbp and large plasmids with genome size $\geq$ 19 kbp. Our results reveal that plasmids in the two size categories rarely share gene families (Fig. SN2.2A). The most shared gene families encode NikA relaxase (in 4% small and 21% large plasmids), Tnp26 (IS*26* transposase; in 2% small and 47% large plasmids) and a Tn*3* resolvase (in 2% small and 19% large plasmids). The absence of gene sharing between small and large plasmids indicates that small and large plasmids are not closely evolutionary related and furthermore shows that there is rarely gene flow between the two groups. Previous studies suggested that small plasmids often co-reside with large plasmids that may also contribute to their mobility[3,4]. To test if gene flow is possible in principle between small and large *KES* plasmids, we tested for co-residence of small and large plasmids in our data. We find that 95% (1,861 out of 1,961) small plasmids reside in the same isolate with at least one large plasmid in all three genera. Taken together, our results show that gene flow between large and small plasmids is limited and restricted to only rare transposition events.

The pattern of shared genes between small and large plasmids reveals two gene families that are highly frequent in plasmids depending on their size: *rop* (*rom*) and *repB* (Fig. SN2.2B). These two proteins are part of two fundamentally different plasmid control or replication mechanisms[5].

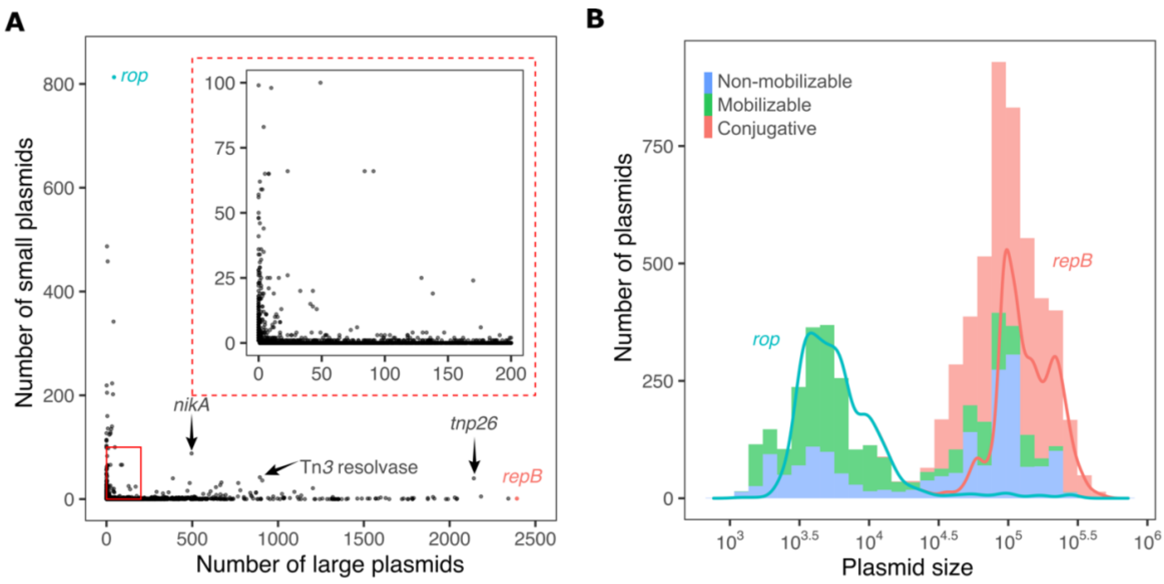

**Figure SN2.2. Identification of marker genes for plasmid size.** (**A**) Shared genes between small (<19Kb) and large (>=19Kb) plasmids. Each dot corresponds to a gene family with the axes corresponding to the number of small and large plasmids harboring a homolog in the family. Most plasmid gene families comprise homologs either in small or large plasmids and are rarely shared between the two size groups. Rop homologs (in blue) are mostly found in small plasmids and RepB homologs (in red) are exclusively found in large plasmids. The three most frequently shared gene families are pointed out and marked with their names. The distribution of gene families present in <100 small plasmids and <200 large plasmids is magnified in the inlay figure. (**B**) A histogram of genome size for all plasmids in the dataset colored by plasmid mobility types. The superimposed density curves (solid line) were created from the size distributions of Rop-encoding plasmids (in blue) and RepB-encoding plasmids (in red).

The *rop* gene family includes 903 homologs in 857 plasmids, of which 95% are smaller than 19 kbp. The *repB* gene family includes 2,603 homologs in 2,393 plasmids. Only one *repB*-encoding plasmid has a genome size smaller than 19 kbp, which we suspect to be a truncated assembly (based on sequence identity to other large plasmids). Taken together, the presence of *rop* or *repB* in plasmid genomes is well associated with plasmid size, with only few exceptions (Fig. SN2.2B; Supplementary Table S3). The distinct gene content of small and large plasmids confirms that plasmid size is associated with different plasmid replication backbones (i.e., plasmid types, replication mechanisms). Plasmids containing *rop* or *repB* correspond to 51% (3,483/6,784) plasmids in the total set, hence additional plasmid backbone genes may be associated with plasmid size (Table below). In the following we compare the distribution of ARGs in small and large plasmids by focusing on Rop-encoding and Rep-encoding plasmids.

To further test the utility of the marker genes, we compared the PTU classification[6] with our classification into small and large plasmids. Considering the PTUs that are represented in our dataset, we find that 63 PTUs have a majority of large plasmids and 26 PTUs have a majority of small plasmids. The large PTUs have on average 99.7% large plasmids and the small PTUs have on average 98.9% small plasmids. Thus, we conclude that PTUs are size specific. The marker genes we identified here are useful for the distinction between small and large plasmids, which largely corresponds also to the plasmid mobility class (see Fig. 3).

Based on our analysis we identified additional 9 gene families that may serve for the distinction between small and large plasmids:

Additional gene families for the distinction of plasmid size types

| | Plasmid type | RefSeq Accession | Gene family |
|---|---|---|---|
| 1 | small | WP_223277951.1 | MbeB family mobilization protein |
| 2 | small | WP_062955149.1 | MobC family plasmid mobilization relaxosome protein |
| 3 | small | WP_021527181.1 | MbeD family mobilization/exclusion protein |
| 4 | small | WP_001271348.1 | replication initiation protein |
| 5 | large | WP_000109071.1 | translesion error-prone DNA polymerase V autoproteolytic subunit |
| 6 | large | WP_000290782.1 | single-stranded DNA-binding protein SSB |
| 7 | large | WP_004146678.1 | plasmid segregation protein ParM |
| 8 | large | WP_001104873.1 | conserved protein with unknown function YubE |
| 9 | large | WP_001568041.1 | DNA methylase YubD |

### Supplementary Tables

**Supplementary Table S1. A list of 9,225 replicons used in searching for coARGs.**
See supplied Excel file SuppTables.

**Supplementary Table S2. ARG families in plasmid resistance islands (REI).**
See supplied Excel file SuppTables. The table is ordered according to the frequency in REI.

**Supplementary Table S3. The frequency of significant cooccurring ARGs (coARGs) in replicons, syntenic blocks and transposable elements.**
See supplied Excel file SuppTables.

**Supplementary Table S4. Biased distribution of transposases and site-specific recombinases (SSRs) in plasmids and chromosomes.**
See supplied Excel file SuppTables. The table is ordered according to the frequency in REI.

**Supplementary Table S5. A list of significantly co-occurring ARGs in ARG-encoding plasmids.**
See supplied Excel file SuppTables. The table comprises three components: K: *Klebsiella* plasmids; E: *Escherichia* plasmids; S: *Salmonella* plasmids.

**Supplementary Table S6. CSBs distribute biased towards specific PTU.**
See supplied Excel file SuppTables.

**Supplementary Table SN1. Example pieces of resistance island.**
See supplied Excel file SuppTables. CSBs are shown in Supplementary Note SN1.

### Supporting Data.

Phylogenetic data of five transposase gene families, Rop, and RepB. For each of the data, homologs protein sequences, protein information, sequence alignment and phylogenetic tree are supplied.

Collinear syntenic blocks (CSB) data, protein family information, and a guide video are available in https://www.mikrobio.uni-kiel.de/de/ag-dagan/ressourcen

### Supplementary Figures

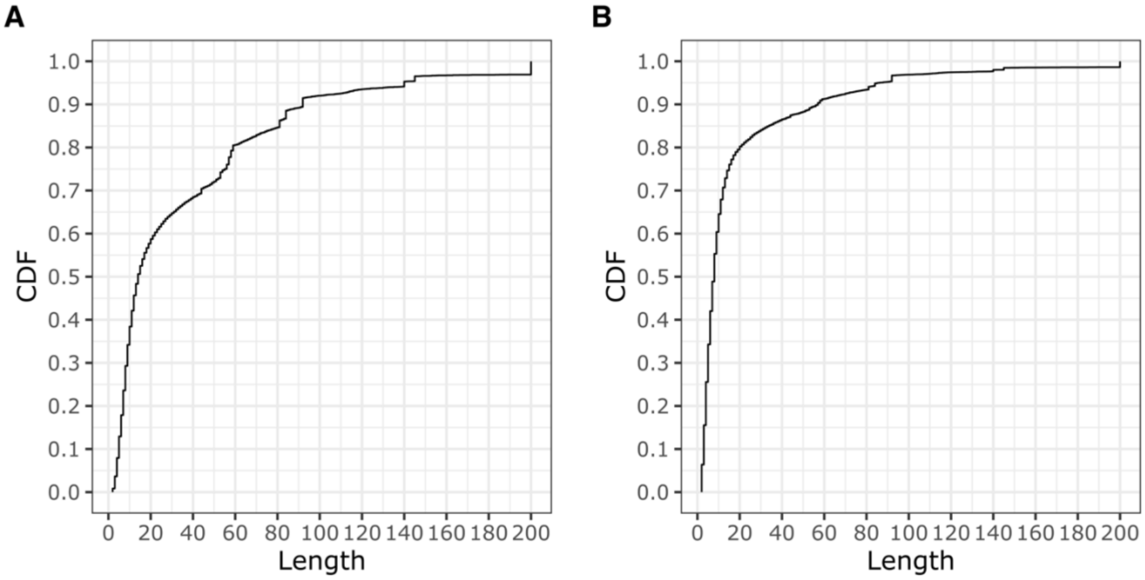

**Supplementary Figure S1. Cumulative distribution function (CDF) of the length of CSBs containing coARGs and matching to transposable elements.**
The CSB length is calculated as the number of protein-coding genes in each CSBs. **A**. A CDF of CSB length for all unique CSBs (i.e., not taking into account the frequency of their occurrence in plasmids). The median length of unique CSBs is 14 genes. **B**. A CDF of all CSB instances in plasmid genomes. The median CSB length across all plasmids and instances is 8 genes.

**A**

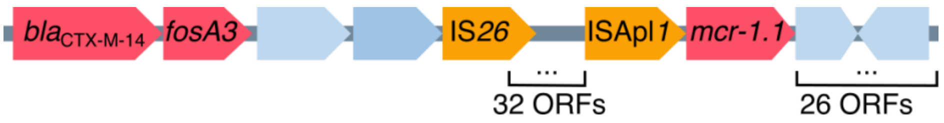

**B**

| Plasmid | RefSeq Accession | Length (bp) | Host | Strain | Isolation date | Isolation location | Isolation source |
|---|---|---|---|---|---|---|---|
| p2474-MCR1 | NZ_CP021209.1 | 223,982 | *E. coli* | Z247 | 2015 | China:Hefei | Blood |
| pTB501 | NZ_CP034820.1 | 188,527 | *Salmonella* sp. | SSDFZ54 | 2018 | China:Zhejiang | Feces |
| pCFSA629 | NZ_CP033351.2 | 210,674 | *S. enterica* | CFSA629 | 2016 | China:Guangdong | Egg |
| pS438 | NZ_CP061125.1 | 253,817 | *S. enterica* | S438 | 2016 | China:Hangzhou | Stool |

**Supplementary Figure S2. A demonstrative example for a CSB corresponding to conserved MGE neighborhood.**

(**A**) A 65-genes long CSB which was found in four plasmids reported in four different genomes. The two ARG-family pairs *fosA3* & *mcr-1.1* and *fosA3* & *bla*$_{CTX\text{-}M\text{-}15}$ were found as significantly cooccurring. The three ARGs are further found within a single CSB, which is constituted of 65 protein-coding-genes including three ARGs. A part of this CSB matches to know transposable elements (e.g., Tn*6330*[7]). (**B**) The CSB was observed in four *Escherichia* and *Salmonella* plasmids. We compared the gene content of the four plasmids and confirmed that they are not identical, yet, all four plasmids include this CSB as a common sequence.

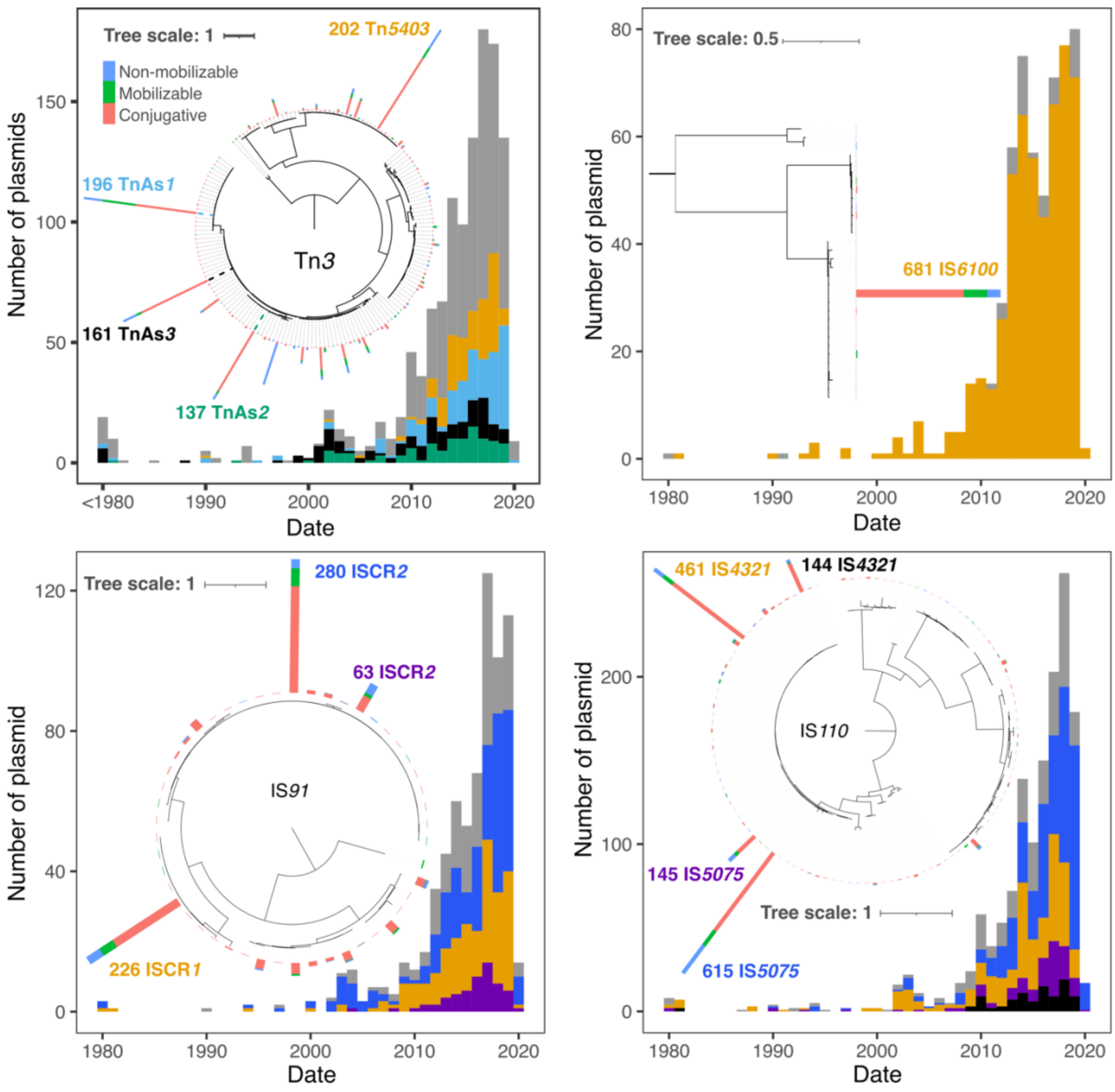

**Supplementary Figure S3. Phylogeny and timeline distribution of SSRs of Tn*3* family, IS*6100*, IS*91*, and IS*110*.**

TnAs*1* (WP_001138014.1), TnAs*2* (WP_001138073.1), and TnAs*3* (WP_001138064.1) belong to the Tn*21* clade in Tn3 family. Tn*5403* (WP_001553819.1, also found in Tn*4378.1* and Tn*511*) belongs to the Tn*163* clade in Tn*3* family. Stacked bars at the tree leaves show the number of the dominant variants, the bar is colored according to the mobility type of the variant carrying plasmid. Histogram below the phylogeny shows the isolation dates of plasmids carrying variants in different shades.

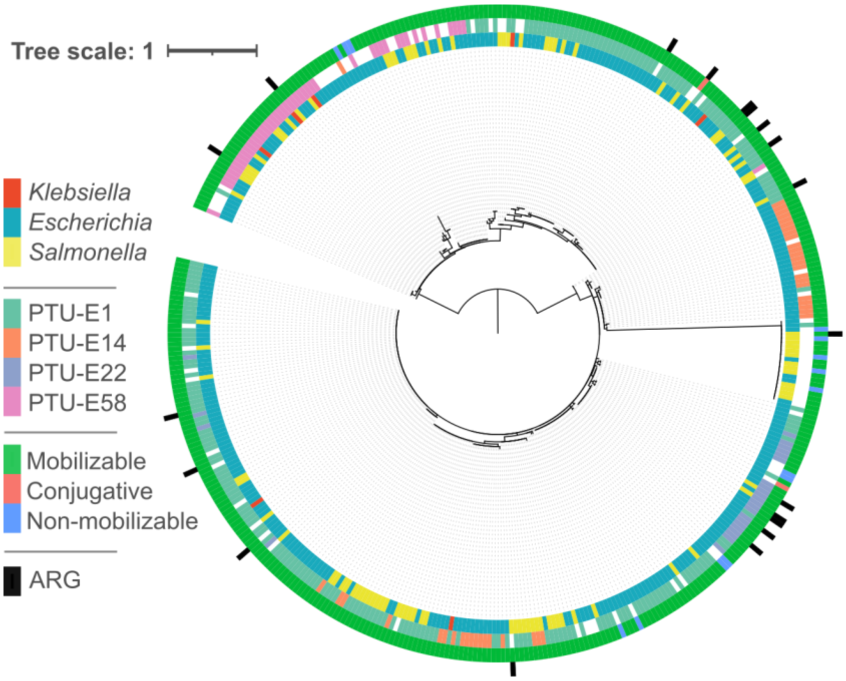

**Supplementary Figure S4. Phylogenic tree of 422 Rop protein sequences in 405 plasmids mostly present in *Escherichia* and *Salmonella*.**
In *Escherichia* and *Salmonella* Rop-plasmids rarely encode ARGs. Rings of the phylogenetic tree show (from inner to outer ring): host genus, plasmid PTU, plasmid mobility type (see legend), and the presence of ARGs.

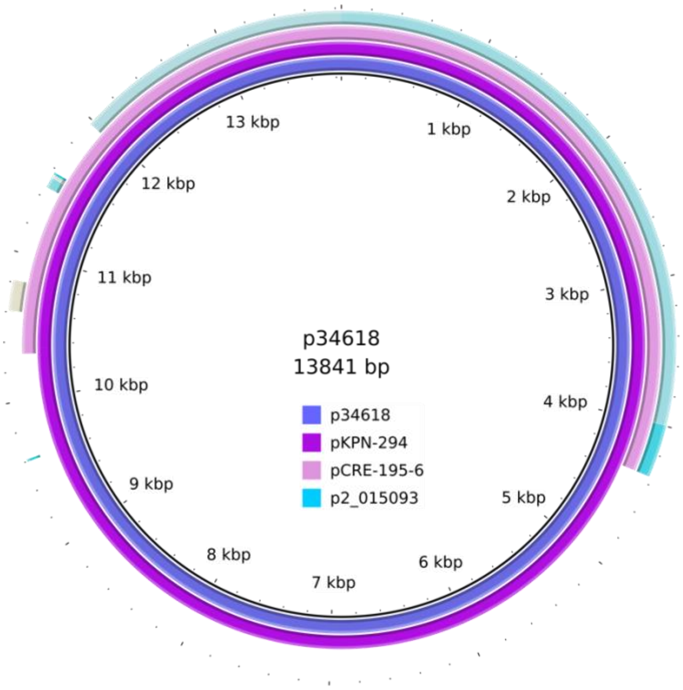

**Supplementary Figure S5. Genome comparison of Rop-encoding plasmids.**
A comparison between four Rop-encoding plasmids selected from the two sister groups of ColRNAI (PTU-E4) mobilizable plasmids (in Fig. 4). ABR plasmids p34618 (NZ_CP010394.1 used as reference genome) and pKPN-294 (NZ_CP009873.1) containing Tn*1331*Δ:IS*26*[5] have the almost identical plasmid backbone as the non-ABR plasmids pCRE-195-6 (NZ_CP061395.1) and p2_015093 (NZ_CP036303.1) from the sister group.

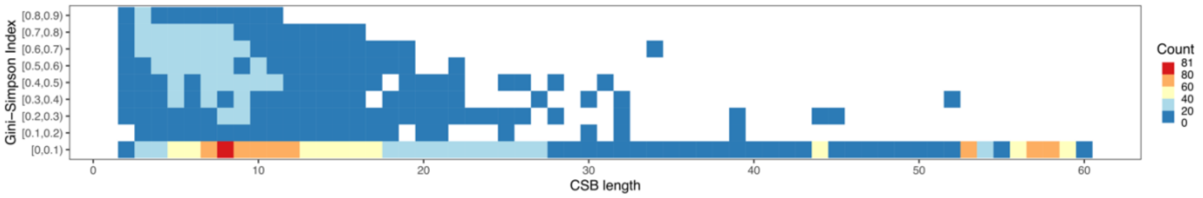

**Supplementary Figure S6. Plasmid diversity and CSB length are associated.** Plasmid diversity was calculated according to the Gini-Simpson Index in 89 identifiable PTUs. Plasmid diversity values were classified into nine groups with interval of 0.1, no value exceeded 0.9. CSBs longer than 60 ORFs (not shown here) have a plasmid diversity index value less than 0.1.

### References

1. Poirel, L. *et al.* Tn*125*-related acquisition of $bla_{NDM}$-like genes in *Acinetobacter baumannii*. *Antimicrob Agents Chemother* **56**, 1087–1089 (2012).
2. Smillie, C., Garcillan-Barcia, M. P., Francia, M. V., Rocha, E. P. C. & de la Cruz, F. Mobility of plasmids. *Microbiol Mol Biol Rev* **74**, 434–452 (2010).
3. San Millan, A., Heilbron, K. & MacLean, R. C. Positive epistasis between co-infecting plasmids promotes plasmid survival in bacterial populations. *ISME J* **8**, 601–612 (2014).
4. Barry, K. E. *et al.* Don't overlook the little guy: An evaluation of the frequency of small plasmids co-conjugating with larger carbapenemase gene containing plasmids. *Plasmid* **103**, 1–8 (2019).
5. Ramirez, M. S., Iriarte, A., Reyes-Lamothe, R., Sherratt, D. J. & Tolmasky, M. E. Small *Klebsiella pneumoniae* plasmids: neglected contributors to antibiotic resistance. *Front Microbiol* **10**, 2182 (2019).
6. Redondo-Salvo, S. *et al.* Pathways for horizontal gene transfer in bacteria revealed by a global map of their plasmids. *Nat Commun* **11**, 3602 (2020).
7. Li, R. *et al.* Genetic characterization of *mcr*-1-bearing plasmids to depict molecular mechanisms underlying dissemination of the colistin resistance determinant. *J Antimicrob Chemother* **72**, 393–401 (2017).
